# Cortical response variability is driven by local excitability changes with somatotopic organization

**DOI:** 10.1101/2022.04.26.489557

**Authors:** T. Stephani, B. Nierula, A. Villringer, F. Eippert, V.V. Nikulin

## Abstract

Identical sensory stimuli can lead to different neural responses depending on the instantaneous brain state. Specifically, neural excitability in sensory areas may shape the brain’s response already from earliest cortical processing onwards. However, whether these dynamics affect a given sensory domain globally or occur on a spatially local level is largely unknown. We studied this in the somatosensory domain of 38 human participants with EEG, presenting stimuli to the median and tibial nerves alternatingly, and testing the co-variation of initial cortical responses in hand and foot areas, as well as their relation to pre-stimulus oscillatory states. We found that amplitude fluctuations of initial cortical responses to hand and foot stimulation – the N20 and P40 components of the somatosensory evoked potential (SEP), respectively – were not related, indicating local excitability changes in primary sensory regions. In addition, effects of pre-stimulus alpha (8-13 Hz) and beta (18-23 Hz) band amplitude on hand-related responses showed a robust somatotopic organization, thus further strengthening the notion of local excitability fluctuations. However, for foot-related responses, the spatial specificity of pre-stimulus effects was less consistent across frequency bands, with beta appearing to be more foot-specific than alpha. Connectivity analyses in source space suggested this to be due to a somatosensory alpha rhythm that is primarily driven by activity in hand regions while beta frequencies may operate in a more hand-region-independent manner. Altogether, our findings suggest spatially distinct excitability dynamics within the primary somatosensory cortex, yet with the caveat that frequency-specific processes in one sub-region may not readily generalize to other sub-regions.

## Introduction

Moment-to-moment fluctuations of neural responses to sensory stimuli play a critical role in how we perceive the external world. Commonly, it is assumed that instantaneous neural states influence the stimulus-related brain responses, which in turn shape the perceptual outcome (e.g., Arieli et al., 1996; McCormick et al., 2020; Sadaghiani et al., 2010; Stephani et al., 2021). Specifically, this may be achieved through the modulation of cortical excitability, which is assumed to shift the sensory threshold of stimulus perception (Samaha et al., 2020). Using electroencephalography (EEG) or magneto-encephalography (MEG), this phenomenon can be observed in various sensory modalities in humans, such as the visual (Busch et al., 2009; Iemi et al., 2017), auditory (Henry et al., 2016; Müller et al., 2013), and somatosensory domain (Baumgarten et al., 2016; Craddock et al., 2017; Forschack et al., 2020; Stephani et al., 2021), where instantaneous neural states have typically been quantified by prestimulus oscillatory activity in the alpha frequency range (8-13 Hz), a common marker of the excitability state of a given cortical brain region (Jensen and Mazaheri, 2010; Klimesch et al., 2007; Romei et al., 2008). Noteworthy, in the somatosensory domain, also activity in the beta frequency range (15–30 Hz) may have a similar modulatory effect on sensory processing (Anderson and Ding, 2011; Jones et al., 2010; van Ede et al., 2011).

Certainly, such fluctuations of cortical excitability – as for example reflected in the dynamics of alpha oscillatory activity – should not be understood as a single, homogenous brain rhythm, but rather as complex network activity involving many distinct neural sources and processes (Buzsáki and Draguhn, 2004; Nikulin et al., 2011; Nunez et al., 2001; van der Meij et al., 2016; Varela et al., 2001). On the one hand, effects of the alpha rhythm on perception have been shown to be brain region-specific (Romei et al., 2008), and spatially fine-tuned by attention even within a sensory modality, reflected for example in retinotopic (Popov et al., 2019) and hand-specific modulations of alpha (and beta) activity (Anderson and Ding, 2011; Jones et al., 2010; van Ede et al., 2011). On the other hand, there are reports of changes in neural states that exert rather unspecific, global effects on perception, for example, associated with a general arousal level (Gee et al., 2020; Schröder et al., 2020), which is also known to be related to alpha oscillations in humans (Barry et al., 2007). Although arousal fluctuations are just one possible explanation for excitability changes, it could thus be that sensory domains are affected by these dynamics as a whole (i.e., a “global effect” within a given sensory domain) or that such global effects even coexist with local, content-specific modulations (Podvalny et al., 2019) that shape perceptual content through spontaneous changes of pre-stimulus activity (Hesselmann et al., 2008). The notion of wide-spread dynamics of cortical excitability is further supported by findings in the motor domain, where moment-to-moment variability of TMS-elicited motor-evoked potentials extended even to the contralateral hemisphere (Ellaway et al., 1998). Given that both sensory and motor processing reflect activity of broad neural circuitries, understanding neural response variability requires a proper specification of where and when these changes in excitability states take place. In this context, it has been found for the somatosensory domain that fluctuations of cortical excitability emerge already at earliest cortical processing in primary sensory regions (Stephani et al., 2020), with behaviorally relevant effects on stimulus intensity perception (Stephani et al., 2021). However, the spatial organization of these neural dynamics is not understood well yet, leaving open the question of whether instantaneous changes of excitability at earliest cortical processing follow local fluctuations or reflect a global state of the primary somatosensory cortex.

Following up on a previously developed technique to probe instantaneous changes of cortical excitability by short-latency somatosensory-evoked potentials (SEP) in the human EEG (Stephani et al., 2021; Stephani et al., 2020), we here examined the trial-to-trial dependencies between neural responses to somatosensory stimulation of hand and foot regions (Fig. 1), as well as their relation to ongoing oscillatory states. Specifically, the N20 component of the SEP, a negative deflection after 20 ms at centro-parietal electrode sites in response to median nerve (i.e., “hand”) stimulation (Allison et al., 1991), is thought to be exclusively generated by excitatory post-synaptic potentials (EPSPs) of the first thalamo-cortical volley (Bruyns-Haylett et al., 2017; Nicholson Peterson et al., 1995; Wikström et al., 1996), and therefore represents a direct measure of instantaneous cortical excitability. Assuming a homologous neural circuitry for the somatosensory foot region, we related this measure to the first cortical component of the SEP in response to tibial nerve (i.e., “foot”) stimulation, the P40 component (Allison et al., 1996; Kany and Treede, 1997). This way, we sought to dissociate local from global fluctuations of neural excitability across spatially distinct sites within the somatosensory cortex. In addition, we compared these dynamics between a long and a short inter-stimulus interval (ISI), as well as in alternating vs. single nerve stimulation to control for influences of the experimental stimulation paradigm. Overall evidence was in favor of a somatotopic organization of excitability effects during early cortical processing, consistent with the idea of local rather than global neural dynamics at the beginning of the neuronal response cascade in the human somatosensory cortex.

**Fig. 1.**
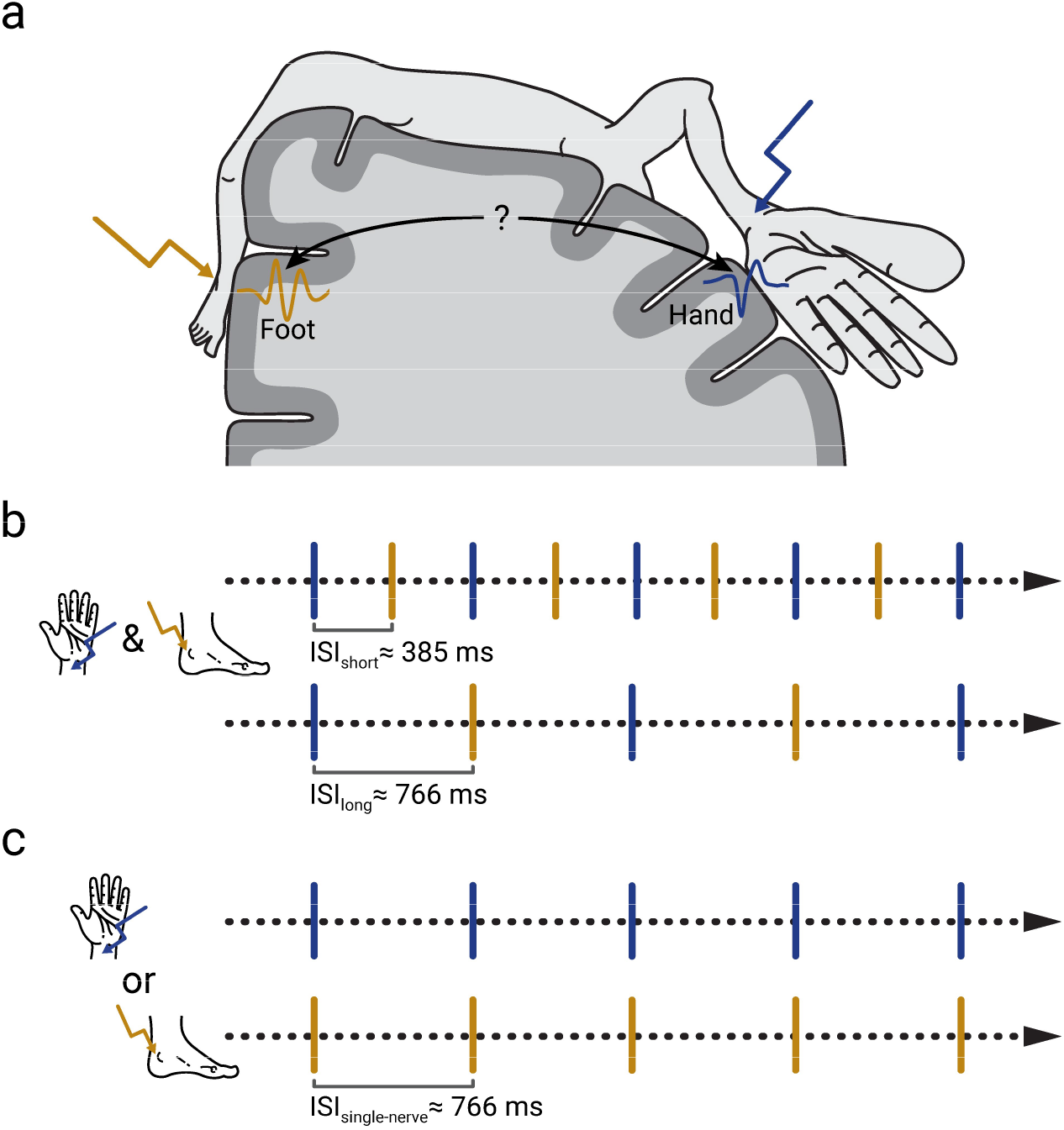
Experimental paradigm. **a)** Schematic of the representation of different body parts in the primary somatosensory cortex. Responses to hand and foot stimuli emerge at spatially distinct sites. **b)** Electrical stimulation of the median (hand) and tibial (foot) nerves was applied in an alternating sequence. In two sub-samples (N=23 & N=15), different inter-stimulus intervals (ISI) were presented: ISI_short_ = 385 ± 50 ms and ISI_long_ = 766 ± 50 ms. **c)** Control conditions of median and tibial stimulation alone (both performed in all participants).

## Methods

### Participants

In total, 42 participants took part in this study, from whom 40 complete datasets could be obtained. Two more participants had to be excluded since no clear SEPs could be extracted on a single-trial level, resulting in a final sample of 38 participants in the alternating stimulation conditions (52.6% female; mean age = 25.7 years, *SD* = 3.7); in the single-nerve control conditions, one additional participant had to be excluded due to similar reasons, leading to a sample of 37 participants. All participants were right-handed as assessed with the Edinburgh Handedness Inventory (Oldfield, 1971; lateralization score, M = + 79.0, SD = 18.2) and did not report any neurological or psychiatric disease. The participants were recruited from the database of the Max Planck Institute for Human Cognitive and Brain Sciences, gave informed consent, and were reimbursed monetarily. The study was approved by the local ethics committee at the Medical Faculty of Leipzig University.

### Stimuli

Somatosensory stimuli were administered by means of electrical stimulation of the median and of the tibial nerve. Non-invasive bipolar stimulation electrodes were positioned on the volar side of the left wrist and on the median side of the left ankle (cathode always proximal), for median and tibial nerve stimulation, respectively. The electrical stimuli consisted of 0.2 ms square-wave pulses and were presented by a DS-7 constant-current stimulator (Digitimer, Hertfordshire, United Kingdom). The intensity of the stimuli was set just above the motor threshold, so that a muscle twitch of the thumb or big toe was clearly visible for every stimulus, with average intensities of 6.77 mA (*SD* = 2.07) for median nerve stimulation and 12.30 mA (*SD* = 3.55) for tibial nerve stimulation. The stimulation was perceived by all participants as a distinct but not painful sensation.

### Procedure

The present study was part of a more extensive project with simultaneous electroencephalo-, electrospino-, electroneuro-, electrocardiographic, and respiratory recordings (Nierula et al., in prep.). After preparation of the electrodes, participants lay down on their backs on a massage bed in a semi-darkened and noise shielded EEG cabin. First, a 5-min resting-state EEG was recorded, followed by 4 blocks of median and 4 blocks of tibial nerve stimulation alone. These single-nerve stimulation conditions served as control conditions for the subsequent alternating stimulation condition. The single-nerve stimulation blocks were presented in an alternating order, counter-balanced across participants, with breaks in between blocks. Each block contained 500 stimuli and the inter-stimulus interval (ISI) was set to 766 ms with a uniformly distributed jitter between +50 and −50 ms. This ISI distribution served to prevent any phase alignment of stimulus onsets with 50 Hz power line artifacts or other periodic noise sources and also ensured the close comparability with a previous study of ours (Stephani et al., 2020). After these blocks, median and tibial nerve stimuli were presented alternatingly within the same blocks (referred to as the alternating stimulation condition). Here we used two different ISIs of ISI_short_ = 385 ± 50 ms and ISI_long_ = 766 ± 50 ms (between subsequent stimuli at alternating sites; both including a uniformly distributed jitter), presented in different sub-samples of participants (N_short_ = 23; N_long_ = 15). The long ISI was chosen in accordance with the single-nerve stimulation condition and the short ISI was set at around half the duration, ensuring that the ISI was still long enough not to lead to any repetition-related attenuation of the initial cortical SEP components (Wikström et al., 1996). In the short-ISI condition, 1000 stimuli each were presented to the median and tibial nerves, while 500 stimuli each were presented in the long-ISI condition. In both cases, the stimuli were split into two blocks, separated by a short break. During the experimental blocks, participants were instructed to look at a fixation cross attached to the ceiling of the cabin. The duration of all experimental blocks was approximately 90 minutes.

Since we were interested in investigating concurrent changes of excitability at different cortical locations, we also considered a simultaneous application of median and tibial stimuli. However, given the spatial overlap between both types of SEPs in the EEG we decided not to perform such stimulation. We rather carefully selected interleaved patterns of alternating stimulation, which on the one hand allowed us to investigate the activation of each cortical area separately and reliably, and on the other hand enabled us to have stimuli sufficiently close in time in order to assess the near-simultaneous processing of both stimuli.

### Data Acquisition

EEG data were recorded from 64 Ag/AgCl electrodes at a sampling rate of 10000 Hz with an 80-channel EEG system (NeurOne Tesla, Bittium, Oulu, Finland), employing a built-in band-pass filter in the frequency range from 0.16 to 2500 Hz. Electrodes were positioned in an elastic cap (EasyCap, Herrsching, Germany) at the international 10-10 system locations Fp1, FPz, Fp2, AF7, AF3, AFz, AF4, AF8, F9, F7, F5, F3, F1, Fz, F2, F4, F6, F8, F10, FT9, FT7, FT8, FT10, FC5, FC3, FC1, FCz, FC2, FC4, FC6, C5, C3, C1, Cz, C2, C4, C6, CP5, CP3, CP1, CPz, CP2, CP4, CP6, T7, T8, TP7, TP8, P7, P5, P3, P1, Pz, P2, P4, P6, P8, PO7, PO3, PO4, PO8, O1, and O2,with the reference placed on the right mastoid process, POz used as the ground, and an additional electrode located on the left mastoid process for offline re-referencing.

In addition, we measured the compound nerve action potentials (CNAP) of the median and tibial nerves to control for peripheral nerve variability. For the median nerve CNAP, two bipolar electrodes were placed on the inner side of the left upper arm along the path of the median nerve, at a distance of about 3 cm. The tibial nerve CNAP was acquired from a patch of 5 electrodes, placed on the backside of the left knee, with the middle electrode on the center of the popliteal fossa, the other 4 electrodes symmetrically arranged around it at a distance of about 1 cm, and referenced to an additional electrode located about 3 cm proximal to the popliteal fossa. The impedances of all electrodes were kept below 10 kΩ.

### EEG pre-processing

The EEG pre-processing pipeline was adapted from two preceding studies with complimentary research foci (Stephani et al., 2021; Stephani et al., 2020). First, stimulation artifacts were cut out and interpolated using Piecewise Cubic Hermite Interpolating Polynomials (MATLAB function *pchip*). The time windows of interpolation were determined on an individual basis depending on the duration of artifact contamination, starting at −1.5 ms and ending between 3 ms and 6 ms relative to stimulus onset. Next, the data were down-sampled to 5000 Hz (including the default anti-aliasing filter of EEGLAB function *pop_resample*). For examining short-latency SEPs, the EEG data were then band-pass filtered between 30 and 200 Hz, sliding a 4^th^ order Butterworth filter forwards and backwards over the data to prevent phase shift (MATLAB function *filtfilt*). On the one hand, this filter served to specifically focus on the N20 and P40 potentials of the SEP, which emerge from frequencies above around 35 Hz, and to omit contributions of later (slower) SEP potentials of no interest. On the other hand, this filter removed slow trends in the data, reaching an attenuation of 30 dB at 14 Hz, thus making sure that variability of the SEP did not arise from fluctuations within slower frequencies (e.g., alpha or theta band activity). At the same time, this high-pass filter obviated the need for an additional baseline correction. Additionally, the data were visually inspected for segments showing muscle or non-biological artifacts, which were excluded from further analyses. After re-referencing to an average reference, eye, heart, and prominent muscle artefacts were removed using independent component analysis (Infomax ICA) whose weights were calculated on the data band-pass filtered between 1 and 45 Hz (4^th^ order Butterworth filter applied forwards and backwards) before. For the SEP analysis, the data were segmented into epochs from −100 to 600 ms relative to stimulus onset, resulting in (average across participants) 1951, 961, 1973, and 1972 trials for the short-ISI, long-ISI, median-only, and tibial-only condition, respectively. EEG pre-processing was performed using EEGLAB (version 14.1.2; Delorme and Makeig, 2004), and custom written scripts in MATLAB (version 2021a; The MathWorks Inc., Natick, Massachusetts).

### Single-trial extraction using CCA

Single-trial SEPs were extracted using Canonical Correlation Analysis (CCA), as proposed by Waterstraat et al. (2015), and applied in the same way as described in Stephani et al. (2020; 2021) for similar datasets. For multi-channel signals ***X*** and ***Y***, CCA finds the spatial filters ***w***_*x*_ and ***w***_*y*_ that maximize the correlation between the two signals:

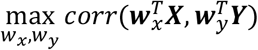

where ***X*** is constructed from concatenating all the epochs of a subject’s recording, i.e. ***X*** = [*x*_1_, *x*_2_, … *x*_*N*_] with *x*_*i*_ ∈ ℝ^*channel × time*^ being the multi-channel signal of a single trial and *N* the total number of trials. In contrast, ***Y*** contains the grand average of all trials, duplicated according to the number of single trials, 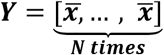 with 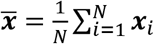. Thus, this CCA procedure can be viewed as a template matching between the single trial signals and a template time signature of the SEP of interest. Solving the optimization problem of CCA (using eigenvalue decomposition) results in a set of weights (i.e., eigenvectors), referred to as the spatial filters ***w***_*x*_, which serve to mix the channels of each single trial (i.e. 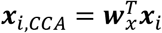) in order to obtain the underlying SEP in CCA space. To focus on the early portion of the SEP only, the two signal matrices ***X*** and ***Y*** were constructed using segments from 5 to 80 ms post-stimulus. The extracted CCA spatial filters were, however, applied to the whole-length epochs from −100 to 600 ms. The signal resulting from mixing the single trial’s channels using the CCA spatial filter ***w***_*x*_, i.e. 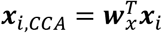, is called a CCA component of that trial *i*.

To obtain the spatial activation pattern of each CCA component we multiplied the spatial filters ***w***_*x*_ by the covariance matrix of ***X***, as *cov*(***X***)***w***_*x*_, thus taking the noise structure of the data into account (Haufe et al., 2014). The resulting spatial activation pattern hence reflects the contribution of each EEG sensor to a given CCA component.

CCA was separately applied to median and tibial stimulation trials individually for every participant. For median stimulation, CCA components whose spatial activity patterns showed the orientation of a tangential dipole over the central sulcus (typical for the N20 potential) were selected for further analyses and referred to as *median CCA components*. In contrast, for tibial stimulation, CCA components were chosen that were characterized by a central radial dipole over the medial part of the primary somatosensory cortex (typical for the P40 potential; referred to as *tibial CCA components*). Such median and tibial CCA components were present for all participants among the first two CCA components with the maximum canonical correlation coefficients, respectively (i.e., one median and one tibial CCA component were selected per participant). Since CCA solutions are insensitive to the polarity of the signal, we standardized the resulting CCA components by multiplying the spatial filter by a sign factor, in the way that the N20 potential in median nerve stimulation always appeared as a negative peak, and the P40 potential in tibial nerve stimulation as a positive peak.

### SEP peak amplitudes and pre-stimulus oscillatory activity

N20 peak amplitudes were defined as the minimum value in single-trial SEPs of the median CCA components ±2 ms around the latency of the N20 in the within-subject average SEP. Similarly, P40 peak amplitudes were defined as the maximum value in single-trial SEPs of the tibial CCA components ±3 ms around the latency of the P40 in the within-subject average SEP.

To infer cortical excitability from oscillatory brain activity, we quantified the amplitudes of pre-stimulus oscillatory activity in the alpha and beta frequency bands. While alpha band activity is a well-established indicator of cortical excitability (Jensen and Mazaheri, 2010; Klimesch et al., 2007; Romei et al., 2008), we also included beta band activity in our analyses since it may serve a similar function particularly in the somatosensory domain (Anderson and Ding, 2011; Jones et al., 2010; van Ede et al., 2011). For the extraction of pre-stimulus alpha and beta band amplitudes, the data were segmented from −200 to −10 ms relative to stimulus onset, mirrored to both sides (symmetric padding; in order to avoid filter-related edge artifacts), and band-pass filtered in the alpha band (8 to 13 Hz), as well as in the beta band (18 to 23 Hz), using a 4^th^ order Butterworth filter (applied forwards and backwards). Subsequently, the amplitude envelope in the given frequency band was computed by taking the absolute value of the analytic signal, using Hilbert transform of the real-valued signal. To derive one pre-stimulus metric per trial, amplitudes were averaged across the pre-stimulus time window and log-transformed for subsequent statistical analyses in order to approximate a normal distribution. This analysis was performed both on the signals obtained from the same spatial CCA filters as for the SEP analysis (corresponding to the median and tibial CCA components) as well as in source space (for details of the reconstruction see section *EEG source reconstruction*). With the CCA filter approach, we sought to focus on similar sources in the pre-stimulus data as are involved in the N20 and P40 generation. Complementarily, the source-space-based approach can be viewed as an SEP-uninformed whole-cortex analysis.

In order to avoid block-related sources of variability in the median- and tibial-only control conditions (such as caused by re-adjustments of stimulation intensities between blocks), we here only included the first block for each condition (i.e., approximately the first 500 trials), whereas in the alternating stimulation condition all available trials were used.

### EEG source reconstruction

Source activity of the EEG was reconstructed using a lead field matrix corresponding to a three-shell boundary element model (BEM) computed with OpenMEEG (Gramfort et al., 2010; Kybic et al., 2005) on the basis of a template brain anatomy (ICBM152; Fonov et al., 2009) with standardized electrode locations. We constrained the lead field matrix to sources perpendicular to the cortex surface (5001 sources), inverted it using the eLORETA method (Pascual-Marqui, 2007), and reconstructed the sources for the spatial patterns of the median and tibial CCA components of every subject, as well as for pre-stimulus alpha and beta activity (on a single-trial level). Brainstorm (Tadel et al., 2011) was used for building the head model and visualizing the data in source space. The MATLAB implementation of the eLORETA algorithm was obtained from the MEG/EEG Toolbox of Hamburg (METH; https://www.uke.de/english/departments-institutes/institutes/neurophysiology-and-pathophysiology/research/research-groups/index.html).

### Processing of peripheral electrophysiological data (median and tibial nerve CNAP)

Analogously to the EEG data, stimulation artifacts in the peripheral electrophysiological data were cut out and interpolated using Piecewise Cubic Hermite Interpolating Polynomials. We chose interpolation windows of −2 to 4 ms and −6 to 6 ms relative to stimulus-onset, for the median and tibial CNAPs, respectively (due to the higher stimulation intensity of the tibial stimuli, larger stimulation artefacts occurred in the tibial CNAP). Furthermore, among the five tibial CNAP channels, the channel with the largest average response was selected individually per participant to be used in further analyses. To achieve a sufficient signal-to-noise ratio (SNR) of the short-latency CNAP peak of only a few milliseconds duration on single-trial level, the data were high-pass filtered at 70 Hz (4^th^ order Butterworth filter applied forwards and backwards). Single-trial peak amplitudes were extracted as the minimum amplitude ±1 ms around the participant-specific latency of the CNAP peak that were found between 5 and 9 ms, as well as between 7 and 12 ms in the within-participant averages, for median and tibial stimuli, respectively. In the analysis of these peripheral nerve activity measures, a sub-sample of 25 participants was included who showed clear responses for both the median and tibial CNAPs (exclusions were mostly due to unclear tibial CNAPs).

### Phase slope index (PSI)

In order to examine interactions within the alpha and beta frequency bands in hand and foot regions, we used the phase slope index (PSI), a connectivity metric that indicates the direction of information flow and that is insensitive to spurious connections due to volume conduction (Nolte et al., 2008), defined as follows:

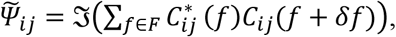

where 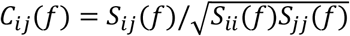 is the complex coherency between two signals *i* and *j*, *S* the cross-spectral matrix, *δf* the frequency resolution, *F* the set of frequencies over which the slope is summed, ∗ indicates the complex conjugate, and ℑ(∙) denotes taking the imaginary part.

For the extraction of the PSI metrics, we first defined regions of interest (ROI) in the hand and foot areas of the right primary somatosensory cortex. The included vertices of the head model were manually selected via the Brainstorm GUI, based on the Destrieux atlas (Destrieux et al., 2010), resulting in 20 and 24 vertices for the hand and foot regions, respectively. After projecting the EEG data into source space (as described above), principal component analysis (PCA) was used to reduce the dimensionality of the ROI activities to one signal trace each (with the largest eigenvalue). These signals were segmented into epochs from – 400 to 400 ms around stimulus onset and the PSI was calculated for frequency bands from 7.5 to 13.75 Hz and from 18.75 to 25.0 Hz for alpha and beta activity, respectively. Slightly wider frequency ranges and a longer time window were chosen here, as compared to the above-mentioned analyses on the pre-stimulus effects, in order to include more frequency bins and to improve the frequency resolution, thus leading to better slope estimates of the phase lags for the PSI calculation. Additionally, pre- and post-stimulus time windows were analyzed separately; please see Supplement D. Finally, PSI values were normalized by their standard deviation as suggested by Nolte et al. (2008):

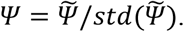

We used the PSI implementation of the MEG/EEG Toolbox of Hamburg (METH; https://www.uke.de/english/departments-institutes/institutes/neurophysiology-and-pathophysiology/research/research-groups/index.html).

### Statistical analyses

The spatial specificity of fluctuations of N20 and P40 potentials was examined using a model-based approach that compared both their within- and across-stimulation-site dependencies over time. For this, we employed cross-lagged two-level structural equation modeling (elsewhere also referred to as *Dynamic Structural Equation Models*; Asparouhov et al., 2018; McNeish and Hamaker, 2020), based on the general latent variable framework of *Mplus* (Muthén and Muthén, 1998-2017). Autoregressive lag-1 relationships were modelled both for N20 and P40 amplitudes on a single-trial level within participants, as well as their cross-lagged dependencies (i.e., the effect of the preceding P40 on the subsequent N20, as well as the N20’s effect on the subsequent P40). Furthermore, in a second approach, the moderating effect of the inter-stimulus interval (ISI) on the inter-relations between N20 and P40 amplitudes was tested as a level-2 covariate. Finally, median and tibial CNAP peak amplitudes were added to the initial model, and both the autoregressive lag-1 effects within these peripheral measures, as well as their relationship with N20 and P40 amplitudes were estimated (this model was calculated in a sub-sample of 25 participants due to the limited availability of clear peripheral CNAPs). In all tested models, path coefficients of the within-participant effects were derived as random slopes using Bayesian model estimation based on Markov-Chain-Monte-Carlo (MCMC) iterations. We used *Mplus*’ default prior distributions (i.e., uninformative priors for path coefficients). The *p*-values provided by *Mplus* correspond to the proportion of coefficients having the opposite sign according to the posterior distribution (reported in order to facilitate a comparison with potentially alternative frequentist statistics). In addition, we indicate the 95%-credible intervals (CI) of path coefficients. Model fit was assessed using the deviance information criterion (DIC; Spiegelhalter et al., 2002), calculated as follows:

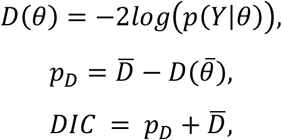

where *θ* represents all model parameters, *Y* all observed variables, *pD* the effective number of parameters (i.e., indicating the model’s complexity; Spiegelhalter et al., 2002), 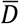 the average deviance, 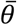 the average model parameters across MCMC iterations (Asparouhov et al., 2018). Please note that traditional fit indices known from maximum-likelihood-basedSEMs(suchas RMSEA, SRMR or CFI) are not available for MCMC model estimation. The DIC can be interpreted in a similar way as the Akaike information criterion (AIC) or the Bayesian information criterion (BIC), that is, lower values indicate a better model fit (Asparouhov et al., 2018; McNeish and Hamaker, 2020). Employing DIC comparisons, the initial cross-lagged SEM was compared to alternative models, stepwise excluding specific effect paths involved in either the cross-lag relationships or the auto-correlations (Table 1). To account for potential instabilities of the DIC values over different MCMC runs (Asparouhov et al., 2018), we repeated the model estimations 100 times each and averaged the resulting pD and DIC values. Subsequently, we tested the differences between models regarding their DIC distributions using pairwise t-tests (*p* values were corrected for multiple comparisons using the Holm-Bonferroni method; Holm, 1979). All physiological measures (i.e., SEPs and CNAPs) included in these models were z-transformed and linear trends were removed before being submitted to the SEM analysis.

**Table 1.** Comparison of alternative SEMs. pD and DIC values were averaged across 100 MCMC runs with different random seeds. Lower DIC values indicate a better model fit. Additionally seeking model parsimony, as reflected in a smaller number of free parameters and lower pD values, SEM 4 presents as the preferred model (marked in bold font).

|  | Model fit indices |  |  |
| --- | --- | --- | --- |
|  | Number of free parameters | Model complexity (effective number of parameters, pD) | Deviance information criterion (DIC) |
| (1) Cross-lagged SEM (incl. auto-regression) | 14 | 1916.3 | 153934.4 |
| (2) Cross-lagged SEM excl. $P40_t \sim N20_t$ | 12 | 1899.3 | 153932.3 |
| (3) Cross-lagged SEM excl. $N20_t \sim P40_{t-1}$ | 12 | 1899.3 | 153936.8 |
| **(4) Auto-regressive SEM excl. cross-lag** | **10** | **1881.2** | **153932.5** |
| (5) Cross-lagged SEM excl. auto-regression | 10 | 1880.8 | 153986.8 |

Effects of pre-stimulus alpha and beta band amplitudes on SEP amplitudes (i.e., N20 and P40) were analyzed using random-intercept linear-mixed-effects models of the following form:

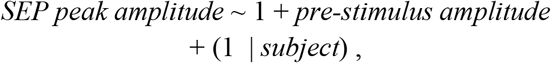

calculated for pre-stimulus amplitudes both in CCA space as well as in source space (mass-univariate approach; i.e., separate models for every source). Furthermore, the moderating effect of the ISI condition was tested by adding an interaction term:

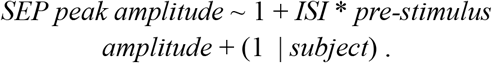

All EEG measures were z-transformed prior to the analysis and pre-stimulus alpha and beta amplitudes were additionally log-transformed in order to approximate a normal distribution. In the mass-univariate approach in source space, we used FDR-correction (*p*<.001) to account for multiple comparisons (Benjamini and Hochberg, 1995). For all analyses (apart from the FDR-correction for multiple comparisons), the statistical significance level was set to *p* = .05.

The linear-mixed-effects models were calculated in R (version 4.1.1, R Core Team, 2018) with the *lmer* function of the *lme4* package (version 1.1-27.1, Bates et al., 2015), estimating the fixed-effect coefficients based on the restricted maximum likelihood (REML). To derive a *p* value for the fixed-effect coefficients, the denominator degrees of freedom were adjusted using Satterthwaite’s method (Satterthwaite, 1946) as implemented in the R package *lmerTest* (version 3.1-3 Kuznetsova et al., 2017). Structural equation modelling was performed in *Mplus* (version 8.6, Base Program and Combination Add-On; Muthén and Muthén, 1998-2017) using the *MplusAutomation* package in R for scripting (version 1.0.0; Hallquist and Wiley, 2018).

### Data availability

The data of the alternating stimulation conditions used in this study are openly available at: https://osf.io/fn7jc/. The data of the single-nerve stimulation conditions can be obtained upon request from the corresponding author (T.S.) and will be published along with an upcoming data article comprising simultaneous electroencephalo-, electrospino-, electroneuro-, electrocardiographic, and respiratory recordings (Nierula et al., in prep.).

### Code availability

The custom-written code that was used for data processing and statistical analyses is publicly available at: https://osf.io/fn7jc/.

## Results

### Short-latency somatosensory evoked potentials on a single-trial level

Single-trial somatosensory evoked potentials (SEPs) were extracted for both the median and tibial nerve stimulation conditions, decomposing the sensor-space EEG by Canonical Correlation Analysis (CCA). As can be seen from Fig. 2, short-latency responses to median nerve stimulation presented with a tangential-dipole pattern with corresponding sources around the hand region of the primary somatosensory cortex, whereas short-latency responses to tibial nerve stimulation were characterized by a radial pattern whose orientation suggested sources in the medial part of the primary somatosensory cortex where the foot region is located. The initial cortical responses of interest, the N20 in median nerve stimulation and the P40 in tibial nerve stimulation, were visible on a single-trial level as negative peak at around 20 ms, and as positive peak at around 40 ms, respectively. Across the stimulation conditions, similar trial-to-trial variabilities were observed regarding N20 and P40 peak amplitudes, with SD_N20, alternating long ISI_ = 0.70 a.u., SD_N20, alternating short ISI_ = 0.78 a.u., SD_N20, single-nerve_ = 0.77 a.u., SD_P40, alternating long ISI_ = 0.58 a.u., SD_P40, alternating short ISI_ = 0.63 a.u., and SD_P40, single-nerve_ = 0.65 a.u. (standard deviations of within-subject trial-to-trial fluctuations, pooled on group level; the units are “arbitrary units”, a.u., due to the normalization of CCA components to have unit variance across the whole training window).

**Fig. 2.**
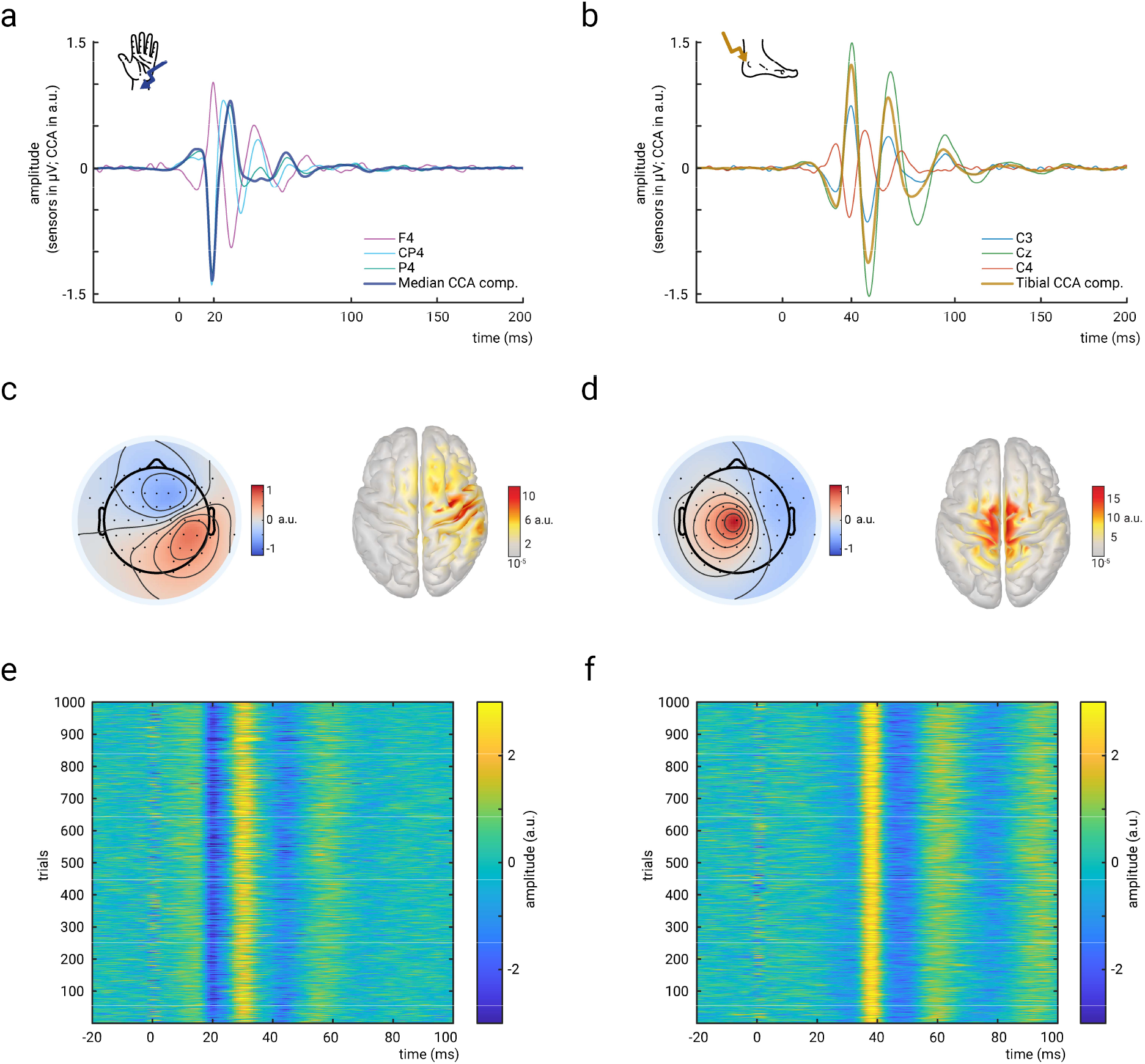
Short-latency somatosensory evoked potentials (SEPs) to median and tibial nerve stimulation. **a)** Grand average of the SEP (N=38) in response to median nerve stimulation, shown for selected electrodes in sensor space, as well as for the representative component of the single-trial extraction using Canonical Correlation Analysis (CCA). **b)** Same as a) but for tibial nerve stimulation. **c)** Spatial activation pattern (left) and source reconstruction (right) of the representative CCA component in median nerve stimulation. The spatial activation pattern indicates the contribution of each EEG sensor to the CCA component activity which was subsequently reconstructed in source space. Averaged across participants (N=38). **d)** Same as c) but for tibial nerve stimulation. **e)** Single-trial responses to median nerve stimulation derived from the representative CCA component of an exemplary participant. **f)** Same as e) but for tibial nerve stimulation. All panels show data from the alternating stimulation condition, pooled across the long- and short-ISI conditions for the group-level averages in panels a-d.

### Short-latency SEP dependencies emerge within but not between stimulation sites

In order to examine whether dynamics in initial cortical somatosensory responses follow local or global dynamics, we tested the inter-dependencies of N20 and P40 peak amplitudes in a cross-lagged two-level structural equation model (SEM). We modelled the lag-1 auto-correlation of subsequent SEPs within hand and foot regions, as well as their cross-lag relationships on a single-trial level. As can be seen from the results of the fitted SEM in Figure 3, only the within-site relationships (i.e., lag-1 auto-regressions) showed significant effects whereas no dependencies emerged between stimulation sites (i.e., cross-lag regressions), *β*_*N20→N20*_ = .039, *p*_*N20→N20*_ < .001, *CI*_*N20→N20*_ = [.024, .055]; *β*_*P40→P40*_ = .021, *p* _*P40→P40*_ = .007, *CI* _*P40→P40*_ = [.004, .038]; *β*_*P40→N20*_ = −.009, *p* _*P40→N20*_ = .146, *CI* _*P40→N20*_ = [−.026, .008]; *β*_*N20→P40*_ = −.006, *p*_*N20→P40*_ = .228, *CI*_*N20→P40*_ = [−.021, .008], respectively (*CI*s correspond to the 95%-credible intervals based on the posterior distributions of the Bayesian model estimation).

**Fig. 3.**
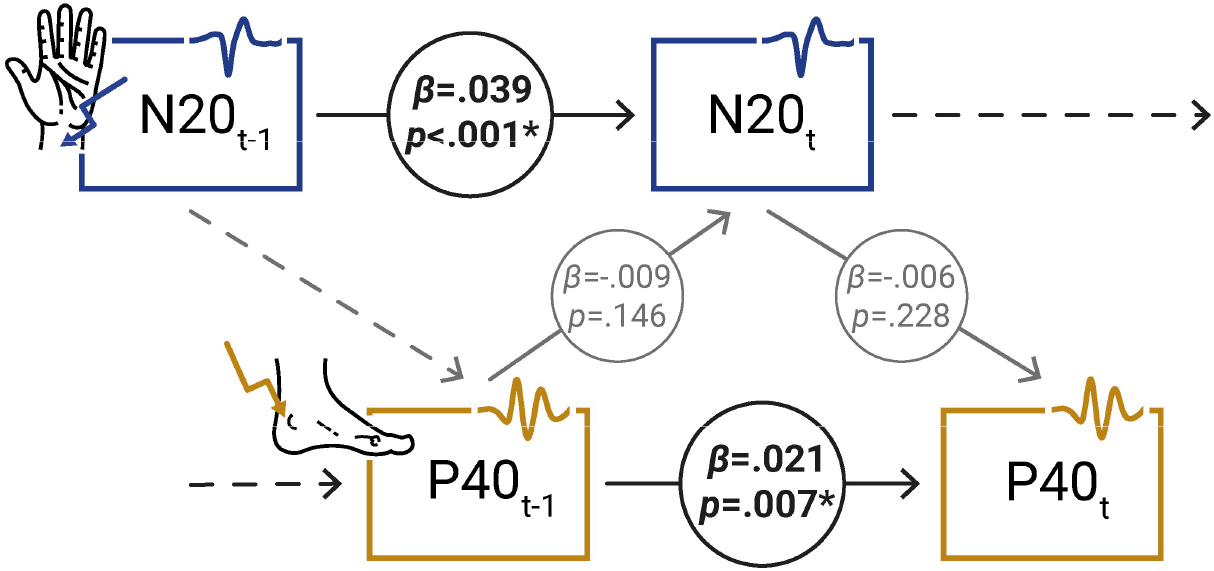
Cross-lagged two-level SEM. Path coefficients reflect mean random slopes on group level, derived from a Bayesian model estimation with one-tailed *p*-values corresponding to the proportion of coefficients having the opposite sign according to the posterior distribution.

These findings were further corroborated by comparisons of alternative models, in which we tested the stepwise exclusion of specific effect paths, that is either paths involved in the cross-lag relationship or in the auto-correlation (Table 1). The models were evaluated using the deviance information criterion (DIC) where lower values indicate better model fit (i.e., in close correspondence to the Akaike information criterion (AIC) in non-Bayesian model estimation). To account for potential instabilities of the DIC values over different MCMC runs (i.e., different model estimation runs) – an issue that has been noted for Bayesian model comparisons before (Asparouhov et al., 2018) – we repeated the model estimations 100 times each and averaged the resulting pD and DIC values. As can be seen from Table 1, SEM 2 and SEM 4 showed the lowest DIC values on average. To confirm this statistically, we examined the differences of DIC values between models using pairwise *t*-tests across the 100 model estimation runs. While SEM2 and SEM4 did not differ, *p* = .839, they both showed significantly lower DIC values than SEM1, SEM3 and SEM5, *p* = .034, *p* < .001, *p* < .001, and *p* = .040, *p* < .001, *p* < .001, for SEM2 and SEM4, respectively (all *p* values Holm-Bonferroni-corrected). Given the equal model fit of SEM2 and SEM4 regarding their DIC values but considering the lower number of model parameters of SEM4 (i.e., seeking model parsimony), we conclude that SEM 4, that is the model without cross-lag relationships, presents as the preferred model to describe the empirical data.

Taken together, both the analysis of path coefficients as well as subsequent model comparisons suggest temporal dependencies of cortical excitability <u>within</u> stimulated regions but not dependencies <u>between</u> the two regions.

In a next step, we added *ISI condition* as an additional between-subjects variable to the original cross-lagged two-level SEM and tested whether the ISI had a moderating effect on either the auto-correlation within cortical regions or on inter-dependencies between the two regions. No moderator effects were observed here, neither for the within-region auto-correlations, *β*_*ISI, N20→N20*_ = −.004, *p*_*ISI, N20→N20*_ = .421, *CI*_*ISI, N20→N20*_ = [−.041, .034], and *β*_*ISI, P40→P40*_ = −.003, *p*_*ISI, P40→P40*_ = .451, *CI*_*ISI, P40→P40*_ = [−.040, .035] (Fig. 4), nor for the cross-lagged relationships between P40 and subsequent N20, as well as N20 and subsequent P40, *β*_*ISI, P40→N20*_ = .003, *p*_*ISI, P40→N20*_ = .446, *CI*_*ISI, P40→N20*_ = [−.034, .039], and *β*_*ISI, N20→P40*_ = −.021, *p*_*ISI, N20→P40*_ = .119, *CI*_*ISI, N20→P40*_ = [−.053, .014], respectively. Thus, different ISIs did not affect the temporal dependencies of short-latency SEP amplitudes, which at the same time endorses our initial approach to pool SEPs from both the long and short ISI (Fig. 2 & Fig. 3).

**Fig. 4.**
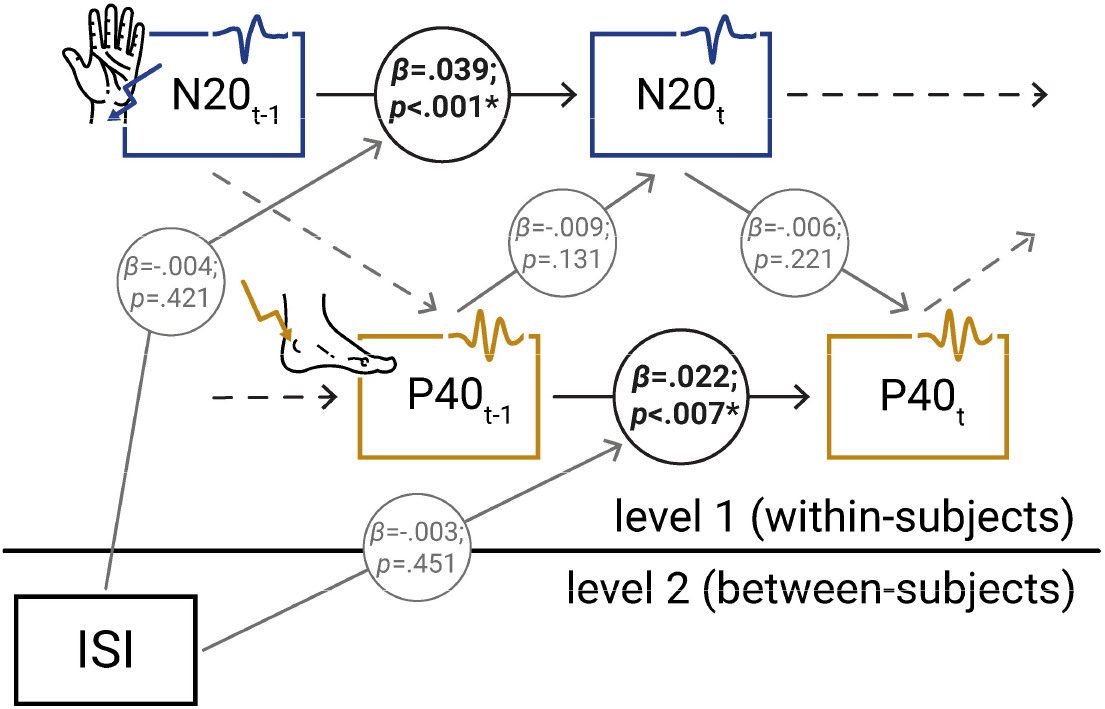
Cross-lagged two-level SEM including ISI as cross-level moderator. Path coefficients reflect mean random slopes on group level, derived from a Bayesian model estimation with one-tailed *p*-values corresponding to the proportion of coefficients having the opposite sign according to the posterior distribution.

Finally, we tested whether the temporal dependencies of SEP amplitudes were driven by changes of peripheral nerve activity, as measured by compound nerve action potentials (CNAPs) at the upper arm and at the back of the knee. In addition to genuine peripheral neural variability, also small displacements of the stimulation electrodes or changes in limb posture could potentially have affected the extent of stimulus-induced nerve excitation and might thus render cortical-excitability explanations for temporal SEP dependencies moot. We employed the initial cross-lagged two-level SEM, but now included the median- and tibial-nerve CNAP measures as additional predictors of N20 and P40 amplitude, respectively. The model results did not indicate any relationship between peripheral nerve activity and SEP amplitudes, *β*_*CNAPmedian→N20*_ = .012, *p*_*CNAPmedian→N20*_ = .309, *CI*_*CNAPmedian→N20*_ = [−.036, .056], and *β*_*CNAPtibial→P40*_ = −.016, *p*_*CNAPtibial→P40*_ = .196, *CI*_*CNAPtibial→P40*_ = [−.060, .024], nor did we observe any auto-correlation within the time-series of CNAP amplitudes, *β*_*CNAPmedian*_ = .033, *p*_*CNAPmedian*_ = .058, *CI*_*CNAPmedian*_ = [−.010, .077], and *β*_*CNAPtibial*_ = −.010, *p*_*CNAPtibial*_ = .338, *CI*_*CNAPtibial*_ = [−.062, .039] – while relations among SEP amplitudes remained comparable (i.e., dependencies within but not between cortical regions; Fig. 5). We therefore conclude that the observed SEP dynamics cannot be explained by peripheral changes of nerve activity.

**Fig. 5.**
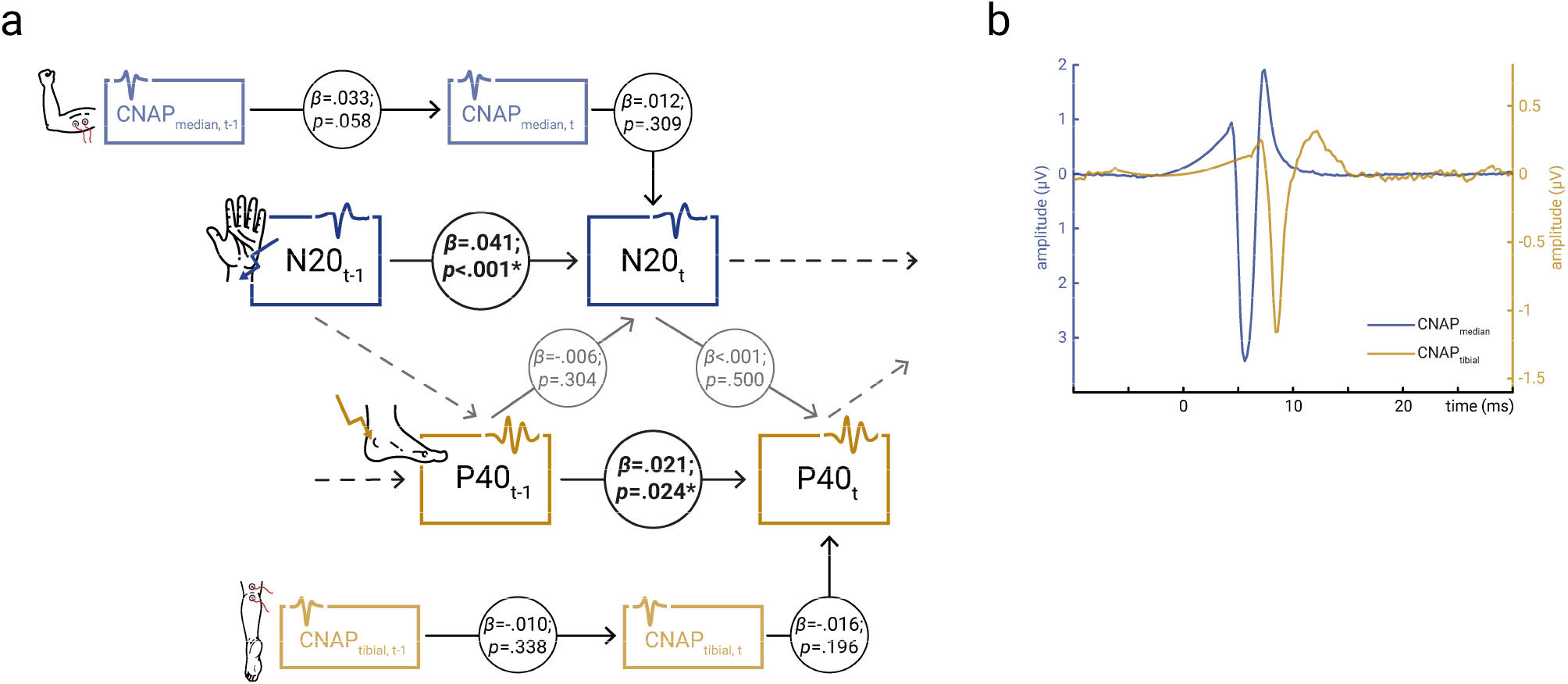
Controls for peripheral nerve variability. **a)** Cross-lagged two-level SEM including peripheral nerve activity measures (*CNAP*_*median*_ and *CNAP*_*tibial*_) as covariates (sub-sample of dataset, N=25). Path coefficients reflect mean random slopes on group level, derived from a Bayesian model estimation with one-tailed *p*-values corresponding to the proportion of coefficients having the opposite sign according to the posterior distribution. **b)** Grand averages (N=25) of the peripheral nerve activity measures *CNAP*_*median*_ and *CNAP*_*tibial*_ in the alternating stimulation condition.

### Spatially confined effects of pre-stimulus alpha oscillatory state on initial SEP amplitudes

Next, we examined the relationship between short-latency SEP amplitudes and pre-stimulus oscillatory state in the alpha band (8-13 Hz). In congruence with previous studies (Stephani et al., 2021; Stephani et al., 2020), we found a negative relationship between the amplitude of the N20 potential (in response to median nerve stimulation) and pre-stimulus alpha amplitude, as extracted with the same spatial CCA filter as the median SEP, *β*_*prestim*_ = −.040, *t*(29611.3) = −6.626, *p* < .001 (calculated with a random-intercept linear-mixed-effects model with N20 amplitude as dependent variable and pre-stimulus alpha amplitude as predictor, pooled across both ISI conditions). Similarly, the P40 potential (in response to tibial nerve stimulation) was related to the pre-stimulus alpha amplitude extracted with the spatial CCA filter of the tibial SEP when pooling both ISI conditions, *β*_*prestim*_ = −.017, *t*(29600.6) = −2.913, *p* = .004. As can be seen from Supplement A, a complementary analysis sorting N20 and P40 amplitudes by quintiles of pre-stimulus alpha amplitudes further supported the linearity of the respective relationships (as opposed to an inverted u-shaped relationship).

In order to test the spatial specificity of the pre-stimulus alpha effects, we repeated the above analyses in source space. For this, we first reconstructed the sources of the pre-stimulus alpha activity and then computed separate linear-mixed-effects models across all sources in the cortex. As shown in Figure 6, the effects of pre-stimulus alpha amplitude on the N20 potential were most prominent in proximity to the hand area in the primary somatosensory cortex, whereas the effects on the P40 potential were located more medially, just above the primary somatosensory foot area. This observation was further supported by the lack of significant pre-stimulus effects for alpha band activity in the primary visual cortex (Supplement C). Thus, in these analyses including both ISI conditions, the effects of pre-stimulus alpha activity appeared to be specific for those spatial regions where the modulated SEP components are generated.

**Fig. 6.**
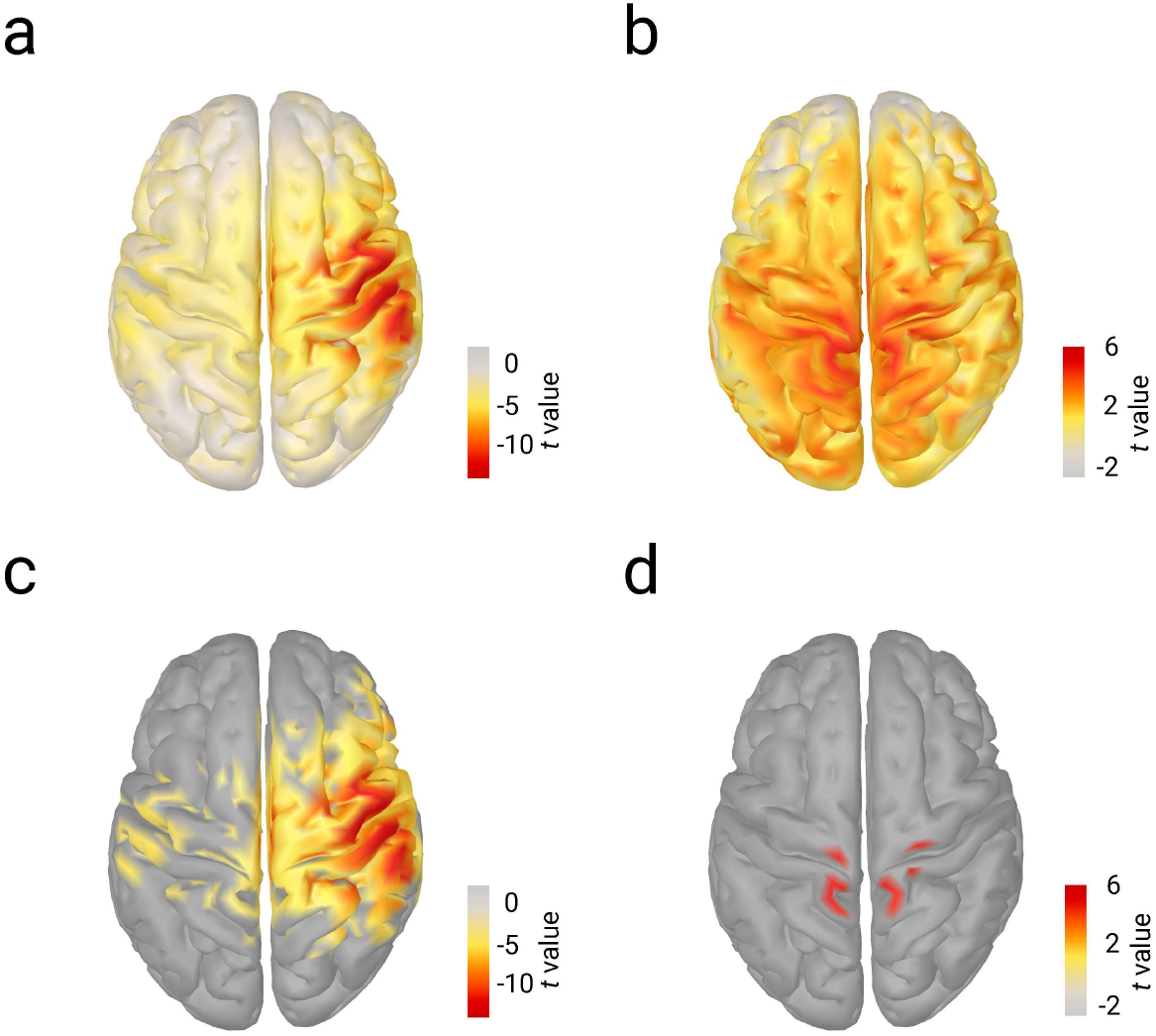
Relationship between pre-stimulus alpha activity and SEPs in source space (pooled ISI conditions). **a)** Effects of pre-stimulus alpha amplitude, reconstructed in source space and averaged between −200 and −10 ms relative to stimulus onset, on N20 amplitude (evoked by median nerve stimulation). Shown are *t*-values corresponding to the *β* coefficients of mass bivariate linear-mixed-effects models in source space (5001 sources; N=38). **b)** Same as a) but for the effect of pre-stimulus alpha amplitude on P40 amplitude (evoked by tibial nerve stimulation). **c)** Same as a) but FDR-corrected to control for multiple comparisons, *p*<.001. **d)** Same as b) but FDR-corrected to control for multiple comparisons, *p*<.001.

### Pre-stimulus alpha effects are consistent across stimulation conditions for N20 but not P40

Although we did not observe any moderating effect of the ISI on the inter-dependencies of short-latency SEP amplitudes (see structural equation models above), the ISI might still play a role for the effect of pre-stimulus alpha activity on SEP amplitudes. To test this, we ran additional control analyses, now including the interaction term *ISI × pre-stimulus alpha amplitude* in the linear-mixed-effects models of the effects of pre-stimulus alpha on N20 and P40 amplitudes, respectively (prestimulus alpha activity was extracted with the same spatial CCA filters as the single-trial SEPs). For the N20 potential, no interaction effect emerged *β*_*ISI×prestim*_ = .006, *t*(29524.0) = .461, *p* = .645, and the main effect of pre-stimulus alpha amplitude remained significant *β*_*prestim*_ = −.045, *t*(29451.6) = −3.697, *p* < .001. Thus, effects of pre-stimulus alpha amplitude on N20 amplitude were comparable for both the long and the short ISI. However, we found a significant interaction effect *ISI × pre-stimulus alpha amplitude* on P40 amplitude, *β*_*ISI×prestim*_ = .033, *t*(29566.9) = 2.460, *p* = .014, and here, the main effect of pre-stimulus alpha on P40 disappeared when including the interaction term, *β*_*prestim*_ = −.008, *t*(29540.1) = −.690, *p* = .490. Further examining this interaction effect with separate linear-mixed-effects models for both ISI conditions revealed that there was an effect of pre-stimulus alpha amplitude on P40 amplitude in the short-ISI condition, *β*_*prestim*_ = .025, *t*(22398.0) = 3.697, *p* < .001, whereas no pre-stimulus effect emerged in the long-ISI condition, *β*_*prestim*_ = −.008, *t*(7193.1) = −.765, *p* = .445. These findings were further supported when examining the pre-stimulus effects in source space but now separately for each ISI condition: While the spatial specificity was consistent for prestimulus alpha effects on the N20, the P40 analyses revealed somatotopic effects in the short-ISI condition only (Fig. 7a).

**Fig. 7.**
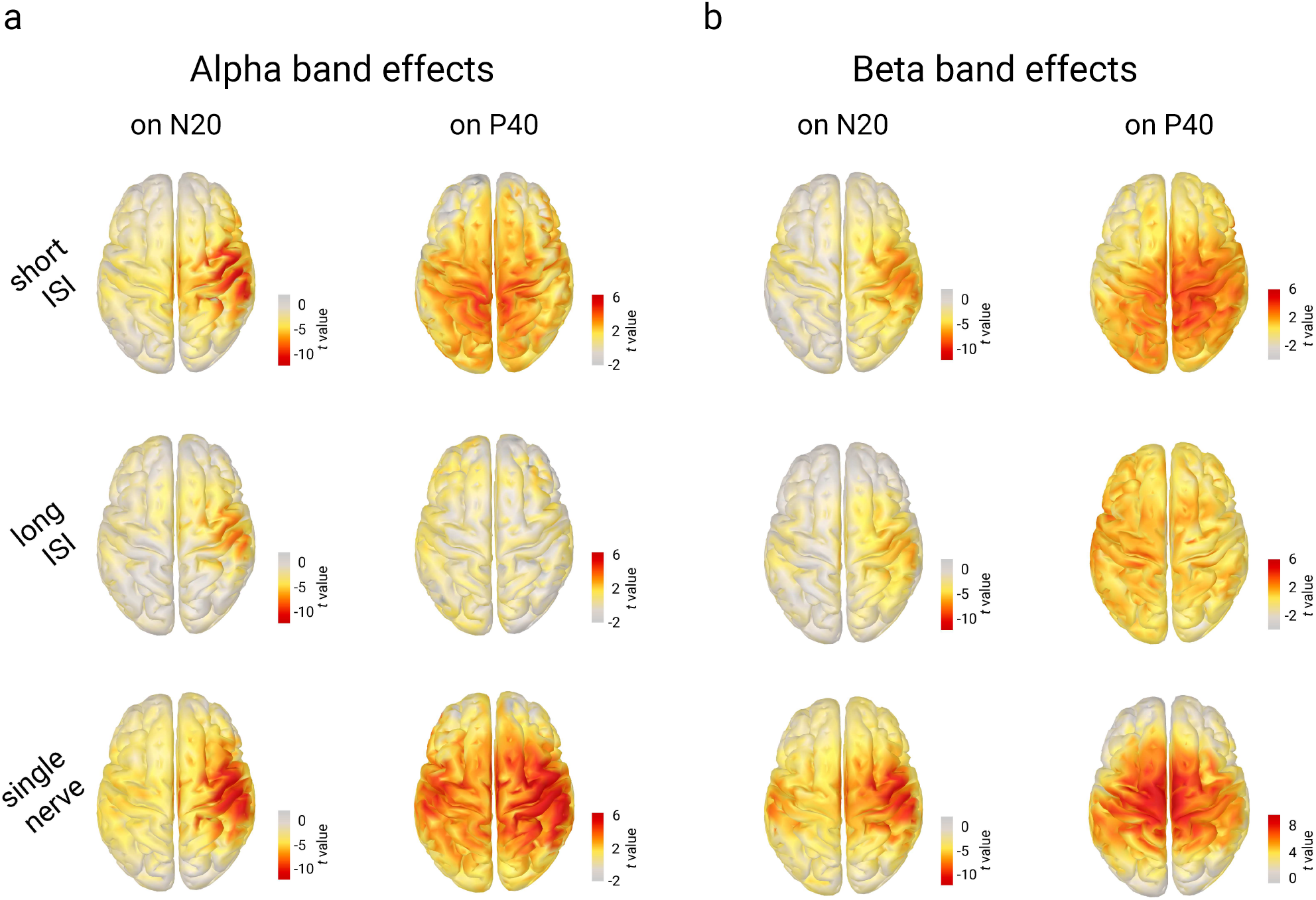
Effects of pre-stimulus alpha (8-13 Hz) and beta (18-23 Hz) band amplitudes on initial SEP amplitudes in source space, displayed across all stimulation conditions. **a)** Alpha band effects on N20 (left column) and P40 (right column), for short ISI (first row; N=23), long ISI (second row; N=15), and single nerve stimulation (third row; N=37). **b)** Beta band effects on N20 (left column) and P40 (right column), for short ISI (first row; N=23), long ISI (second row; N=15), and single nerve stimulation (third row; N=37). All panels display *t*-values corresponding to the *β* coefficients of the effect between pre-stimulus alpha amplitude and N20 or P40 amplitude as calculated by mass bivariate linear-mixed-effects models in source space (5001 sources). Please note the different scaling of the color bar for the beta band effect on P40 in the single nerve condition.

In order to better understand the lack of prestimulus alpha effects on P40 amplitudes in the long-ISI condition (e.g., whether these were specific to the considerably smaller sub-sample of participants we tested here), we re-examined the pre-stimulus alpha effects in additional control conditions in which either median or tibial nerve stimuli were presented alone (ISI ≈ 766 ms). While the relationship between pre-stimulus alpha and N20 amplitudes remained qualitatively comparable with the other median nerve stimulation conditions (Fig. 7a, bottom row left), an unexpected pattern emerged for tibial-nerve-only stimulation: Prestimulus alpha effects on P40 amplitude were most pronounced over bilateral somatosensory hand regions rather than over foot regions (Fig. 7a, bottom row right).

### Spatial specificity of pre-stimulus effects in the beta frequency range (18-23 Hz)

Following up on the heterogenous spatial specificity of pre-stimulus alpha effects in tibial nerve stimulation, we explored whether differential patterns would be observed for the second prominent frequency component of the sensorimotor mu rhythm, the beta frequency band (here measured between 18 and 23 Hz). As pointed out by previous studies that a somatosensory alpha rhythm may be more difficult to detect in foot regions (Pfurtscheller et al., 1997) and given that time-frequency analyses suggested additional prestimulus effects in the beta frequency range both in a recent, similar study (Stephani et al., 2021) as well as in the present data (Supplement B), the beta band may thus provide complimentary insights into the frequency-specific spatial organization of excitability effects. As displayed in Figure 7b, the linear-mixed-effects models in source space indeed showed somatotopic effects of pre-stimulus beta amplitude on N20 amplitude in all stimulation conditions, as well as somatotopic effects of prestimulus beta amplitude on P40 amplitude at least in the short-ISI and single-nerve conditions. Thus, an asymmetric pattern was observed for prestimulus effects on P40 amplitudes particularly in the single-nerve stimulation condition: While prestimulus effects were rather located over (bilateral) hand regions in the alpha frequency range (Fig. 7a, bottom row right), pre-stimulus beta effects emerged primarily over foot regions (Fig. 7b, bottom row right). In contrast, the spatial organization of pre-stimulus effects on the N20 was similar both in the alpha and beta frequency range (please refer to Table 2 for a systematic overview).

**Table 2.** Overview of the spatial specificity of pre-stimulus effects of pre-stimulus alpha and beta activity on N20 and P40 amplitudes. Green color indicates findings in line with a local and somatotopic organization of excitability dynamics (whereas null findings are marked in grey and contradictory findings in red).

|  |  | Pre-stimulus effects |  |  |  |
| --- | --- | --- | --- | --- | --- |
|  |  | Alpha band |  | Beta band |  |
|  |  | on N20 | on P40 | on N20 | on P40 |
| Alternating stimulation | Short ISI | hand area | foot area | hand area | foot area |
|  | Long ISI | hand area | no clear effects | hand area | no clear effects |
| Single-nerve stimulation (control condition) |  | hand area | hand areas (bilaterally) | hand area | foot area |

### Alpha and beta bands express different directional connectivity between hand and foot areas

Motivated by some discrepancies of prestimulus alpha and beta effects across stimulation conditions (Table 2), we tried to further disentangle the general regional specificity of these frequency bands by examining their directed connectivity between hand and foot regions. For this, we employed the phase slope index (PSI; Nolte et al., 2008), a measure whose sign indicates the direction of information flow between two cortical areas. We calculated the PSI between a hand and a foot ROI in the right somatosensory cortex and compared the resulting connectivity metrics across stimulation conditions and the two frequency bands, with positive values reflecting that activity in the hand region led activity in the foot region and vice versa for negative values. Since we were interested in overall, that is, prevailing connectivity differences of the frequency bands, we here examined the whole time range from −400 to 400 ms relative to stimulus onset (please refer to Supplement D for separate analyses of the pre- and post-stimulus time windows). As can be seen in Figure 8, alpha band activity was associated with positive PSI values across all stimulation conditions whereas beta band activity showed PSI values around zero and in a slightly negative regime. This suggests that the alpha rhythm in the hand region led alpha activity in the foot region while this tendency was not present (or even slightly reversed) for the beta rhythms. This observation was statistically confirmed using a repeated-measurement ANOVA with the factors *stimulation site*, *frequency band*, and *single-nerve vs. alternating stimulation* (corresponding to Fig. 8a), which showed a main effect of *frequency band*, *F*(1,37) = 22.528, *p* < .001, *η*^2^ = .127 and an interaction effect of *frequency band* by *stimulation site*, *F*(1,37) = 4.517, *p* = .040, *η*^2^ = .010. Also, a second ANOVA confirmed the main effect of *frequency band* when splitting up the alternating stimulation condition into short- and long-ISI conditions (corresponding to Fig. 8b), *F*(1,36) = 26.347, *p* < .001, *η*^2^ = .224 (here, *ISI condition* was treated as between-subject factor while the factor *single-nerve vs. alternating stimulation* in the former analysis was implemented as within-subject factor). Similar relationships between the frequency band and the PSI values were also obtained when considering the pre- or post-stimulus time windows separately (Supplement D). Notably, these connectivity effects cannot be explained by volume conduction since the chosen connectivity metric, the phase slope index, eliminates these spurious relations (Nolte et al., 2008). Next, one might wonder whether the frequency-specific positive and negative PSI values significantly differed from zero, respectively, which would indicate a certain direction of information flow (and not just a difference between conditions). Examining the 95%-confidence intervals (as shown in Fig. 8), it becomes evident that PSI values in the alpha frequency band were all higher than zero while the PSI values in the beta frequency band were significantly negative in tibial nerve stimulation only.

**Fig. 8.**
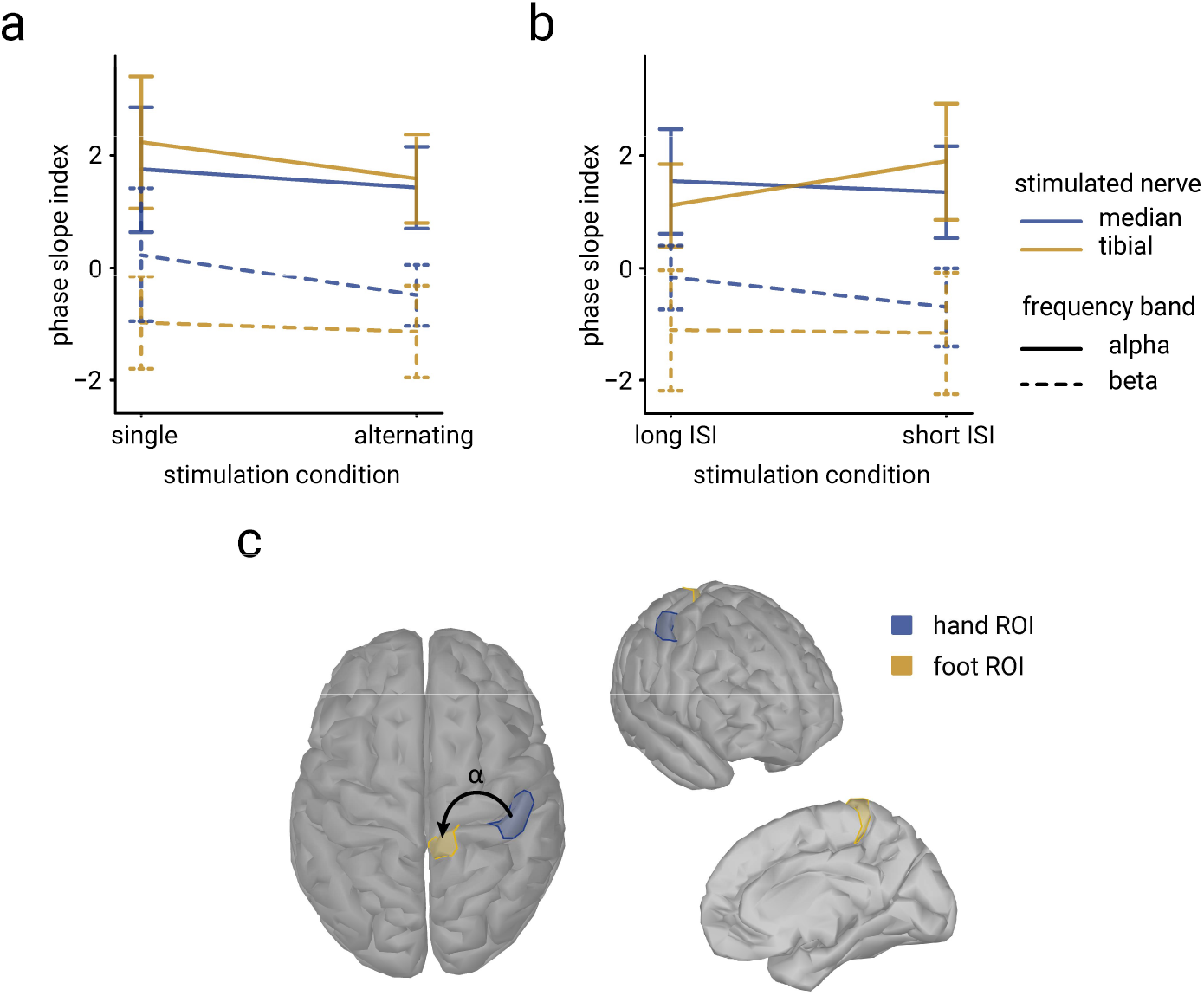
Directed connectivity between hand and foot areas in alpha and beta frequency bands. **a)** PSI values for single nerve vs. alternating nerve stimulation conditions (N=38). **b)** PSI values for long-vs. short-ISI conditions (N=15 and N=23, respectively). **c)** Display of the hand and foot ROIs between which the connectivity metrics were computed, shown from different angles. The arrow in the top view (left cortex surface) illustrates the direction of information flow within the alpha band. PSI values were extracted between −400 and 400 ms relative to stimulus onset. Positive PSI values indicate that activity in the hand region led activity in the foot region and vice versa for negative values. Error bars correspond to the 95%-confidence intervals.

Altogether, these findings thus indicate that alpha band activity reflects a rhythm that is generally orchestrated by activity in the hand region while beta activity does not show this dependency in median nerve stimulation and even tends to be more foot-region-driven in tibial nerve stimulation. At least partially, this asymmetric connectivity pattern may thus explain the divergence of pre-stimulus alpha and beta effects on SEP amplitudes in tibial-nerve-only stimulation.

## Discussion

Combining short-latency somatosensory evoked potentials and pre-stimulus oscillatory state measures, we examined whether instantaneous changes of cortical excitability follow spatially local or global dynamics within the somatosensory domain. We found that initial cortical EEG responses in hand and foot areas showed temporal dependencies regarding their amplitudes, yet only within and not across cortical regions, suggesting local fluctuations of cortical excitability at the earliest processing stages. Furthermore, the relationship between pre-stimulus alpha state and short-latency SEPs was characterized by a somatotopic organization across all hand stimulation conditions. However, this pattern was not equally consistent for foot stimulation where instead beta band activity was found to be region-specific. Offering an explanation for these slightly different effects across regions and frequency bands, post-hoc analyses of directed connectivity suggested that the somatosensory alpha rhythm may be dominated by activity from hand areas while this pattern was not present for beta frequencies.

### Evidence for local excitability fluctuations within primary somatosensory areas

Employing cross-lagged structural equation modeling, we examined the inter-dependencies between short-latency SEPs in the primary somatosensory hand area (N20, evoked by median nerve stimulation) and in the primary somatosensory foot area (P40, evoked by tibial nerve stimulation). According to findings of previous work that the initial component of the cortical somatosensory evoked potential reflects excitatory post-synaptic potentials only (Bruyns-Haylett et al., 2017; Nicholson Peterson et al., 1995; Wikström et al., 1996), N20 and P40 amplitudes represent probes of the instantaneous excitability in the respective somatosensory region at a given time. Our analyses indicated temporal dependencies of instantaneous excitability within but not between primary somatosensory hand and foot regions. These effects occurred irrespective of the ISI conditions – speaking for their robustness – and could furthermore not be explained by variability of peripheral nerve activity. Hence, local fluctuations of neural excitability seem to determine the brain’s response variability. These findings are also well in line with previous work studying the temporal structure of short-latency somatosensory evoked potentials in median nerve stimulation only, where we found that long-range temporal dependencies emerged for even longer time lags within the primary somatosensory hand area (Stephani et al., 2020). Notably, our probing approach using somatosensory stimuli only allows for the observation of excitability fluctuations that occurred on the same (or larger) time scale as the stimulation events (i.e., in the range of seconds), raising the possibility that fluctuations on a smaller time scale might have been missed. We deem this to be unlikely however, since brain responses to stimulations of the *same* body parts were temporally further apart than those of *different* body parts and we did not observe relations between hand and foot region excitability. In addition, fluctuations of cortical excitability in sensorimotor areas, manifested in amplitude modulations of alpha oscillations, are typically characterized by a robust 1/f spectral profile (Linkenkaer-Hansen et al., 2001; Nikulin et al., 2012), which in turn indicates that very slow fluctuations dominate excitability dynamics in the frequency domain (< 0.2 Hz). Therefore, we believe that the ISI(s) used in the present study can be regarded adequate for probing corresponding temporal dynamics.

In a second approach to test the spatial specificity of excitability fluctuations, we related initial cortical stimulus-evoked responses to the pre-stimulus oscillatory state, examined in source space across the cortical surface. In line with the hypothesis that alpha band activity (8-13 Hz) reflects the excitability state of sensory regions (Jensen and Mazaheri, 2010; Klimesch et al., 2007; Romei et al., 2008; Samaha et al., 2020) and replicating the results of a recent study using median nerve stimulation only (Stephani et al., 2021), we observed effects of pre-stimulus alpha amplitudes and initial SEP amplitudes on a single-trial level and now extended these findings also to the foot domain. Although the positive effect direction (i.e., larger alpha amplitudes corresponded to larger SEP amplitudes) may seem counterintuitive at first, it may be explained by the sensitivity of EEG to post-synaptic currents rather than post-synaptic potentials (please refer to Stephani et al. (2021) for a detailed discussion) and also conforms with previous biophysically realistic models of the relation between oscillatory activity and evoked responses in the somatosensory cortex (Jones et al., 2009). Crucially, pre-stimulus alpha effects on evoked potential amplitudes were most pronounced over the primary somatosensory hand area for the N20 and over the primary somatosensory foot area for the P40, thus presenting evidence for a local, somatotopic organization of excitability dynamics. This notion agrees well with recent studies in the visual domain where attention allocation was found to modulate alpha activity with a retinotopic pattern (Popov et al., 2019), associated with changes of the local excitability state (van Kempen et al., 2020), and may reflect the modulation of local neuronal firing rates in the hand area of the primary somatosensory cortex (Haegens et al., 2011).

Notably, the current data supported linear relationships between pre-stimulus alpha amplitude and N20 as well as P40 amplitudes (Supplement A). This may suggest that the local excitability effects we observed here are in fact distinct from previously reported findings on global states of pupil-linked arousal that is typically linked to alpha band power via a non-linear inverted u-shaped relationship (Pfeffer et al., 2022; Podvalny et al., 2021).

### Differences in oscillatory state response dynamics between hand and foot regions

Interestingly though, while the spatial specificity of pre-stimulus alpha effects was consistent across all median nerve stimulation conditions (i.e., most pronounced effects over the stimulated hand region), heterogenous effect patterns emerged across tibial nerve stimulation conditions during control analyses: In the short-ISI condition, pre-stimulus alpha effects were found over the foot region, yet such effects were absent for long ISIs, and in tibial nerve stimulation alone, the most pronounced effects of pre-stimulus alpha amplitude on P40 potentials were instead observed over (bilateral) hand areas (Table 2). This observation appears at odds with the hypothesis of exclusively local, somatotopic dynamics of cortical excitability. On the one hand, the lack of effects in the long-ISI condition may be partially attributable to the smaller number of participants and trials in this condition. On the other hand, the foot area in the primary somatosensory cortex is located much deeper in the brain (in the interhemispheric fissure) and with it the generators of the foot-related P40 as well as of foot-related oscillatory activity are more distant from the EEG sensors as compared to the rather superficial hand area. Thus, it may be more difficult to obtain and correctly localize neural activity from such deeper sources as was reported for foot areas already earlier (Jones et al., 2010). Additionally, the orientation of dipoles in the hand and foot regions should differ due to the cortical folding specific for these regions (which is why the N20 is visible as a negative and the P40 as a positive peak at somatosensory electrodes in the EEG, corresponding to tangential and radial CCA activation patterns, respectively). Although the foot SEPs could be measured very clearly in the current study due to the high stimulus intensity, the dipole orientation difference may still have added to the challenge to observe pre-stimulus narrow-band activity and thus its effects on tibial SEPs. Nevertheless, pre-stimulus alpha effects over the hand regions in the tibial-nerve-only condition in fact rather speak for non-local interactions in the alpha frequency range between foot and hand regions. These differential pre-stimulus dependencies may have at least two reasons. First, it is well known that the somatosensory alpha rhythm is most prominent in hand areas and less consistently found in foot areas (Pfurtscheller et al., 1997). This may be related to the recent finding that different body parts are represented by distinct resting-state functional networks (Thomas et al., 2021), hence possibly also affecting oscillatory rhythms that are thought to arise from activity of neural feedback loops (Bollimunta et al., 2011; Halgren et al., 2019; van Kerkoerle et al., 2014). Second, the segregation of hand and foot areas in the primary somatosensory cortex may be less strict than commonly assumed on the basis of the somatosensory homunculus, with sometimes overlapping representations of distant body parts (Catani, 2017; Muret et al., 2022). In congruence with this account, event-related synchronization of beta band activity after toe stimulation has also been reported in hand regions in humans (in addition to foot regions; Gaetz and Cheyne, 2006) and chemogenetic silencing of hand regions in monkeys may be associated with a disinhibition of foot areas (as measured by BOLD responses using fMRI; Hirabayashi et al., 2021). Thus, more complex inter-areal interactions may occur already at a basic processing level between hand- and foot-related activity, possibly obscuring somatotopic effects of pre-stimulus oscillations.

In another approach, attempting to account for possible differences of hand and foot oscillatory networks, we extended our analyses of prestimulus alpha activity (8 to 13 Hz) to the beta frequency band (18 to 23 Hz), the second major frequency component of the somatosensory mu rhythm, which has been associated with a similar (yet not identical) modulatory role as alpha activity in somatosensory perception (Anderson and Ding, 2011; Jones et al., 2010; Law et al., 2021; van Ede et al., 2010). Here, we indeed found somatotopic effects of pre-stimulus amplitude on P40 amplitude also in the tibial-only condition over more medial regions, presumably corresponding to foot regions, thus contrasting the spatial effect patterns over hand regions in the alpha frequency band. On the one hand, this finding may be related to the assumption that faster oscillatory rhythms (in this case beta) typically originate from neuronal ensembles of smaller sizes (Pfurtscheller and Lopes da Silva, 1999) and might therefore correspond to more local activity than slower rhythms (e.g., alpha). On the other hand, however, this notion cannot entirely explain the asymmetry of effects in alpha and beta bands between the hand and foot stimulation conditions (especially the absence of both alpha and beta effects on P40 amplitudes in the long-ISI condition). Taken together, it appears that somatosensory alpha and beta rhythms in hand and foot regions do not show identical functional properties and that relationships between pre-stimulus states and responses are not readily transferrable across different body regions.

### Hand-dominance of the somatosensory alpha rhythm

Are oscillatory rhythms in hand and foot regions – hypothesized to be indicative of the current neuronal state – independent from each other or do they interact? Addressing this complementary perspective regarding the local versus global organization of excitability within the somatosensory domain, we analyzed the directed connectivity between the two areas in the alpha and beta frequency bands. Strikingly, our analyses identified hand activity to lead foot activity within the alpha but not in the beta band. This suggests that the somatosensory alpha rhythm – also in foot areas – is generally dominated by activity from the hand areas, which may further explain the frequency-specific pre-stimulus effect patterns discussed in the previous section: Since the alpha rhythm in the hand area appears to be a driver for alpha activity in the foot area, both areas are likely to follow similar dynamics within the alpha band. Consequently, since the alpha rhythm is generally most dominant in the hand area, stronger prestimulus alpha effects on foot-related SEP amplitudes are observed over hand rather than foot areas. This adds the important notion that, at least for alpha frequencies, oscillatory rhythms of the somatosensory system may also exhibit components that reflect global coordination, possibly adjusted preferably to the requirements of the hand regions. This is in line with earlier work on the Rolandic mu rhythm over hand regions arguing for both local *and* global alpha activity components (Andrew and Pfurtscheller, 1997) as well as the absence of a fine-grained somatotopy of post-stimulus alpha event-related desynchronization (ERD) on the level of individual fingers (Nierula et al., 2013). Although these asymmetries in somatosensory rhythms between hand and foot areas may seem unexpected, they can be understood from the perspective that hand- and foot-related neural networks serve completely different functions in daily life: While hand coordination often reflects more complex, consciously performed motor sequences (for example in tool use), foot movements are comprised of more monotonous elements and may be executed rather automatically (e.g., when walking). Consequently, these different motoric requirements may also lead to a differential organization of associated sensory functions.

### Conclusions and perspectives

Summing up all findings from the present study, we conclude that overall evidence speaks for modulations of cortical excitability on a rather local than global level – especially when focusing on feedforward sensory processing, such as reflected in N20 and P40 SEP amplitudes. Also, when examining pre-stimulus oscillatory states as markers of excitability, the majority of our findings pointed towards a somatotopic organization of excitability, yet with the limitation that alpha and beta frequency bands may not entirely behave in the same way in hand and foot regions. These inhomogeneities may be due to an asymmetric connectivity pattern of somatosensory oscillatory networks, with the hand region having a prominent role in the alpha frequency range. Despite these presumably local excitability dynamics at early sensory processing, it is conceivable that more global influences come into play at later stages of perception (which we have not focused on here), for example, exerted by top-down processes. Future work may additionally extend the experimental setting to stimuli presented to both sides of the body. This way, it could be studied whether homologous regions in the left and right primary somatosensory cortex (e.g., left- and right-hand areas) share more variance of neural activity than heterologous regions within one hemisphere (e.g., left-hand and left-foot areas), taking into account the existence of interhemispheric connections particularly between homologous S1 regions (Hlushchuk and Hari, 2006; Ragert et al., 2011). Moreover, to further scrutinize the potential role of preceding stimulation events on subsequent pre-stimulus states and evoked responses, a wider range of ISIs should be examined in follow-up studies, possibly also employing less regular (and thus less predictable) stimuli. In this context, it would be interesting to study the specific role of expectation for local excitability dynamics in S1 subregions, given the previous reports of reduced neural gain in case of unpredictable auditory stimuli (Auksztulewicz et al., 2019) and long-term prior effects on the perception of ambiguous visual stimuli (Hardstone et al., 2021). A further question may be whether the spatial specificity of excitability modulations is stable over time or whether the system can adjust it according to the specific task conditions, for example requiring differential involvement of hand and foot neural networks. Last but not least, a promising next step could be to examine the relationship between the here observed local fluctuations of somatosensory excitability in the context of brain-wide, general arousal levels. To this end, future studies could thus combine dedicated arousal measures, such as pupil diameter or skin conductance, with the local excitability metrics employed in the current study, and ideally also include a behavioral control or manipulation of current arousal and/or excitability states.

## Supporting information

Supplement_Stephani et al_Somatotopic_excitability_v2

## Acknowledgements

We are grateful to Janek Haschke, Pia-Lena Baisch, Paula Kosel, Max Braune, Samuel Simeon, Marleen Löffler, and Merve Kaptan for their invaluable help in data collection and their contributions to data pre-processing. Furthermore, we thank Heike Schmidt-Duderstedt for her support in creating Figure 1. This work was funded by the Max Planck Society.

## Author contributions

T.S., B.N., A.V., F.E., and V.V.N. designed research; B.N. and T.S. performed research; T.S. and B.N. analyzed data; T.S. wrote the first draft of the paper; T.S., B.N., F.E., and V.V.N. wrote the paper; T.S., B.N., A.V., F.E., and V.V.N. edited the paper.

