## Supplement_Stephani et al_Somatotopic_excitability_v2 for "Cortical response variability is driven by local excitability changes with somatotopic organization"

### *Supplement A: Binning analysis of pre-stimulus alpha and short-latency SEP amplitudes*

To assess the linearity assumption implied in the linear regression models that were calculated for the effects of pre-stimulus alpha amplitude on N20 and P40 amplitudes, respectively, additional binning analyses were performed. For this, N20 (P40) amplitudes were sorted according to quintiles of pre-stimulus alpha amplitude (Supplementary Fig. 1). Pre-stimulus alpha activity was extracted using the same spatial filters as for the SEP, corresponding to the median and tibial CCA filters for N20 and P40 amplitudes, respectively. Both the relationships of pre-stimulus alpha with N20 as well as with P40 amplitudes appear to be best described by a linear function (as opposed to an inverted u-shaped relationship).

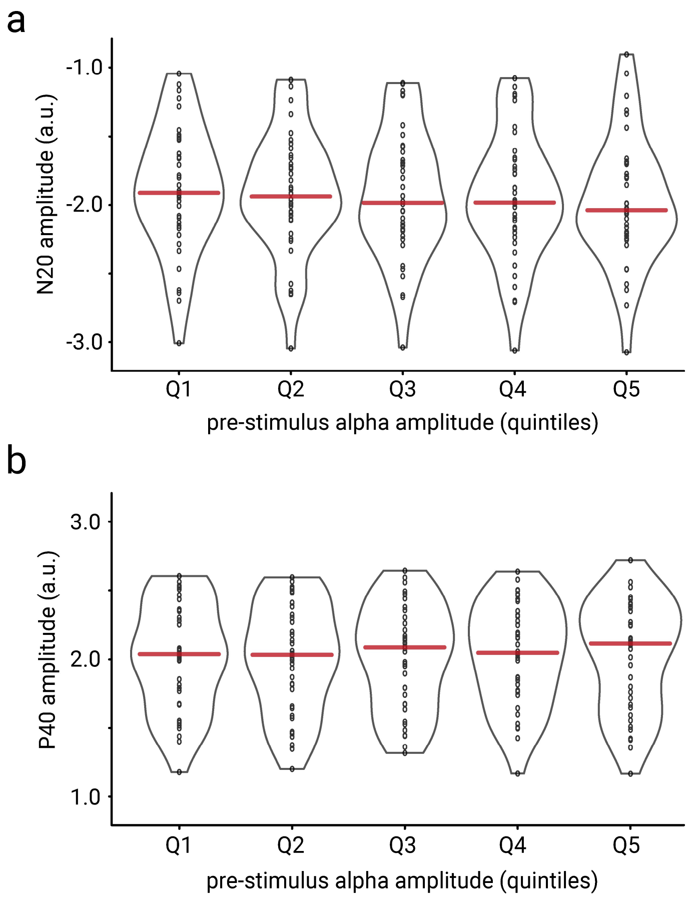


**Supplementary Fig. 1.** Binning analyses of pre-stimulus alpha and short-latency SEP amplitudes. **a)** Relationship between pre-stimulus alpha amplitude and N20 amplitudes. **b)** Relationship between pre-stimulus alpha amplitude and P40 amplitudes. Horizontal red lines indicate the median of the respective distributions. To preserve interpretability of the magnitude of N20 and P40 amplitudes, unstandardized CCA values are shown here (in contrast to the statistical analyses reported in the main manuscript).

*Supplement B: Time-frequency representation of pre-stimulus effects*

In addition, we examined the effects of pre-stimulus alpha activity on SEP dynamics in the time-frequency domain. For this, we again decomposed the multi-channel EEG activity with the spatial CCA filters corresponding to the single-trial SEP components and computed random-intercept linear-mixed-effects models for the amplitude at every frequency and time point regarding its relation to N20 and P40 peak amplitude, respectively:

$$\textit{SEP amplitude} \sim 1 + \textit{t-f amplitude} + (1|\textit{subject})\ .$$

The time-frequency decomposition was performed using complex Morlet wavelets from 4 to 40 Hz (3 to 10 cycles, logarithmically scaled) and time-frequency amplitudes correspond here to the "total" activity, that is, both evoked and induced activity.

Accounting for the observation that pre-stimulus effects on P40 amplitudes were absent in the sample with long ISIs (see section *Pre-stimulus alpha effects are consistent across stimulation conditions for N20 but not P40* in the main manuscript), we restricted the time-frequency analysis for the P40 to the short ISI condition only, whereas for the N20 component, we again pooled the data of both ISI conditions. As can be seen from Supplementary Figure 2, effects of pre-stimulus activity on SEP amplitudes were most prominent in the alpha (around 10 Hz) and beta (around 20 Hz) frequency bands, both for the N20 and P40 analysis. We thus conclude that the link between local pre-stimulus dynamics and SEP fluctuations may indeed be specific to these frequency bands which are indicative for changes of cortical excitability.

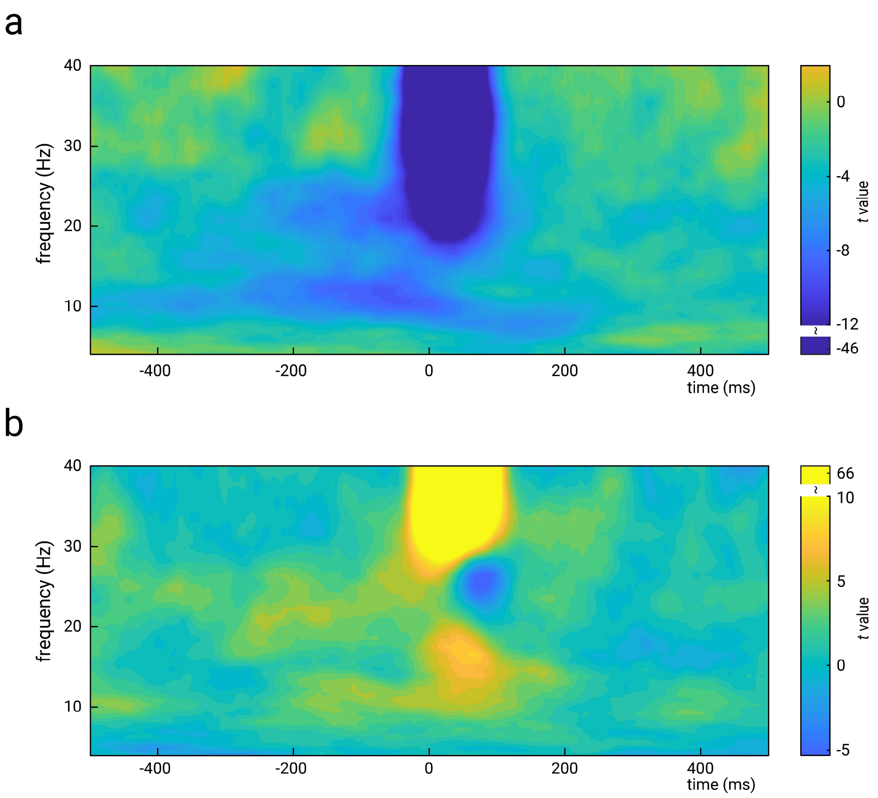


**Supplementary Fig. 2.** Relationship between signal amplitude in the time-frequency domain and SEP amplitudes. **a)** Effects on N20 amplitude (median nerve stimulation), pooled across both ISI conditions (N=38). **b)** Effects on P40 amplitude (tibial nerve stimulation), short-ISI condition only (N=23). Both panels show unthresholded *t*-values of the effect of predictor *t-f amplitude* on *N20*/*P40 amplitude*. Time-frequency amplitudes reflect the "total" activity, that is, both evoked and induced activity, which explains the strong effects for higher frequencies around the expected peak latencies of the N20 and P40 potential, respectively. Please note the discontinued color bars for values below t = -12 in panel a and above t = 10 in panel b (for visualization of the *t*-value ranges of interest).

*Supplement C: No pre-stimulus alpha band effects in the primary visual cortex*

To double-check the spatial specificity of pre-stimulus alpha band effects on SEP amplitudes, we performed a control analysis deliberately focusing on alpha band activity in the primary visual cortex (as opposed to hand and foot regions of the primary somatosensory cortex): In close correspondence to the analyses in the main manuscript, we reconstructed pre-stimulus alpha activity in source space. Next, we extracted the mean pre-stimulus alpha amplitude from the right primary visual cortex (V1), by aggregating pre-stimulus alpha amplitude within the "occipital pole" ROI of the Destrieux atlas (Destrieux et al., 2010). These V1 pre-stimulus amplitudes were now used as predictors of N20 and P40 amplitudes, respectively, employing random-intercept linear-mixed-effect models of the following form:

$$SEP\ amplitude \sim 1 + V1\ alpha\ amplitude + (1|subject)\ .$$

Neither for N20 nor for P40 amplitudes, V1 alpha amplitude effects reached significance, $\beta_{N20} = -.0132$, $t(29442.3) = -1.812$, $p = .070$ and $\beta_{P40} = .0128$, $t(29450.0) = 1.794$, $p = .073$. This further corroborates the notion of local excitability dynamics rather than effects of general brain-wide arousal levels. (The small and non-significant trends in these analyses may have been caused by remaining leakage from somatosensory regions associated with the extensive volume conduction in EEG recordings.)

*Supplement D: PSI analyses in pre- and post-stimulus time windows*

Complementing the analyses of directed connectivity presented in the main manuscript, we also calculated the phase slope index (PSI; Nolte et al., 2008) separately in pre- and post-stimulus time windows, that is, from -400 to -10 ms and 10 to 400 ms, respectively. Again, we examined the PSI between a hand and a foot ROI in the right somatosensory cortex and compared the two frequency bands (alpha and beta), as well as the stimulation conditions by ANOVAs of the same form as in the main manuscript (i.e., with factors *stimulation site*, *frequency band*, *single-nerve vs. alternating stimulation* (repeated-measurement), as well as *ISI condition* (between-participants)).

For the pre-stimulus time window, we found main effects of *frequency band* both in the ANOVA comparing single-nerve vs. alternating stimulation, as well as in the ANOVA comparing long- vs. short-ISI conditions, $F(1,37) = 17.164$, $p < .001$, $\eta^2 = .102$, and $F(1,36) = 21.338$, $p < .001$, $\eta^2 = .224$, respectively. Similarly, main effects of *frequency band* were also found in the post-stimulus time window, $F(1,37) = 30.630$, $p < .001$, $\eta^2 = .206$, and $F(1,36) = 20.219$, $p < .001$, $\eta^2 = .230$ (again for the ANOVAs either comparing single-nerve vs. alternating stimulation or long- vs. short-ISI conditions, respectively). Additionally, a somewhat smaller but significant main effect of *stimulation site* was found in the long- vs. short-ISI ANOVA, $F(1,36) = 6.000$, $p = .019$, $\eta^2 = .015$.

Altogether, these findings match with the results presented in the main manuscript: PSI values are higher (i.e., hand region activity leads foot region activity) for the alpha as compared to the beta frequency band. This seems to be the case across all stimulation conditions and no matter whether the pre- or post-stimulus is analyzed. Therefore, the dominance of the hand region regarding the somatosensory alpha rhythm may be a stable phenomenon, not changed by median and tibial nerve stimulation events.
